# Mu and beta power effects of fast response trait double dissociate during precue and movement execution in the sensorimotor cortex

**DOI:** 10.1101/2024.11.11.621252

**Authors:** Sara Kamali, Fabiano Baroni, Pablo Varona

## Abstract

A better understanding of the neural and muscular mechanisms underlying motor responses is essential for advancing neurorehabilitation protocols, brain-computer interfaces (BCI), feature engineering for biosignal classification algorithms, and identifying biomarkers of disease and performance enhancement strategies. In this study, we examined the neuromuscular dynamics of healthy individuals during a sequential finger-pinching task, focusing on the relationships between cortical oscillations and muscle activity in simultaneous electroencephalography (EEG) and electromyography (EMG) recordings. We contrasted two pairs of subsets of the dataset based on the latency of EMG onset: an across-subjects trait-based comparison and a within-subjects state-based comparison. Trait-based analyses showed that fast responders had higher baseline beta power, indicating stronger motor inhibition and efficient resetting of motor networks, and greater mu desynchronization during movement, reflecting enhanced motor cortex activation. Visual association areas also displayed more pronounced changes in different phases of the task in subjects with lower latency. Fast responders exhibited lower baseline EMG activity and stronger EMG power during movement initiation, showing effective motor inhibition and rapid muscle activation. State-based analyses revealed no significant EEG differences between fast and slow trials, while EMG differences were only detected after movement onset. These results highlight that fast response trait is related to electrophysiological differences at specific frequency bands and task phases, offering insights for enhancing motor function in rehabilitation, biomarker identification and BCI applications.

**Highlights:**

- We analyzed simultaneous EEG and EMG recordings during a sequential finger movement task
- We identified neural correlates of response latency trait, but not state
- Fast responders show higher baseline beta power, indicating stronger motor inhibition
- Lower latency trait was linked to a sharper postcue sensorimotor mu power decrease
- Fast responders have lower pre-movement EMG and higher EMG onset power

## 1. Introduction

Finger movement tasks, such as finger-pinching and finger-tapping, have played an important role in neuroscience and neurorehabilitation research due to their ability to reveal intricate details about motor control mechanisms and motor function in healthy individuals and clinical populations [1–3]. These tasks require precise and coordinated movements, making them excellent paradigms for understanding the interactions between various brain regions [4, 5]. The neural circuitry involved in such tasks includes the primary motor cortex, premotor cortex, sensory regions, basal ganglia, and cerebellum, all of which contribute to the planning, execution, and refinement of motor actions [6–12]. These simple motor tasks have thus served as sensitive indicators of motor system integrity, playing a critical role in the study of both voluntary movement in healthy subjects and its pathological alterations in disease.

The neural substrates of voluntary movement are particularly well studied through finger movement tasks, as they can demonstrate both fine motor control and gross coordination, depending on the nature of the task [13–17]. For instance, finger-pinching tasks, which require opposition of the thumb and one or more fingers, engage fine motor skills and dexterity. These tasks are sensitive measures of motor performance and are often used to evaluate motor deficits in conditions such as Parkinson’s disease, stroke, dystonia, and multiple sclerosis (MS) [18–20]. In MS patients, impairments in fine motor control can significantly affect daily functioning, as even small deficits in upper limb dexterity and coordination can lead to difficulties in performing essential daily activities. Finger-pinching tasks allow clinicians and researchers to assess the extent of these deficits and monitor their progression [21].

Similarly, finger-tapping tasks that involve repetitive tapping movements are utilized to assess motor speed, rhythmicity, and coordination. These tasks have been instrumental in detecting motor abnormalities and monitoring disease progression in neurological disorders such as Parkinson’s disease and MS [22, 23]. For instance, in Parkinson’s disease, bradykinesia (slowness of movement) is often assessed by evaluating the speed and rhythmicity of finger tapping. Motor fluctuations can also be detected through changes in tapping behavior over time [18], making these tasks highly valuable for tracking disease progression and the effectiveness of interventions such as medication or deep brain stimulation.

Evaluative finger movement tasks extend beyond traditional clinical assessments and enter the realm of advanced neurotechnologies, particularly in brain-computer interface (BCI) research [24, 25]. BCI enables direct communication between the brain and external devices. Finger movement tasks are often paradigms to decode motor intentions from neural signals. In BCI research, tasks such as finger-tapping are employed to train machine learning algorithms to recognize motor imagery patterns, facilitating the development of assistive technologies for individuals with motor impairments [26]. These technologies range from prosthetic control systems to neurorehabilitation devices for restoring lost motor function [27–29]. For example, finger-tapping tasks have been used to develop BCIs that restore hand function in patients who have experienced motor deficits due to stroke or spinal cord injury [30–32]. Progress in EEG feature extraction methods improves the classification accuracy of brain signals [33–36], which has application in personalized prosthetic control systems and personalized rehabilitation. These advancements demonstrate the potential of combining finger movement tasks with cutting-edge neurotechnologies to improve rehabilitation outcomes and offer new avenues for improving motor function.

Another critical component of finger movement tasks is the measurement of movement latency—the time between a stimulus (e.g., a visual or auditory cue) and the initiation of movement. Latency provides insights into the efficiency of sensorimotor processing [37]. It can indicate how quickly the brain can translate sensory information into motor action. Shorter latencies are associated with faster neural processing and motor execution. Conversely, prolonged latencies may indicate motor impairments or neurological dysfunctions, making latency measurements valuable in clinical assessments [38]. Analysis of EMG onset latency in healthy subjects during finger-pinching tasks provides insights into the dynamics of neural activity associated with motor execution and control [39].

In healthy individuals, shorter latencies reflect efficient neural processing within the sensorimotor system, but in clinical populations such as Parkinson’s disease, MS, or stroke, increased latency often correlates with disease severity, i.e., prolonged movement latencies due to deficits in motor initiation and execution [40]. These prolonged latencies can result from disruptions in the normal functioning of the basal ganglia, which play a crucial role in initiating and modulating voluntary movements [41]. Understanding the neural and muscular factors contributing to variations in movement latency is crucial for developing targeted interventions to improve motor function in these populations.

In particular, EEG power in the mu and beta bands is highly informative in finger movement studies, providing crucial insights into motor control mechanisms in both healthy and clinical individuals. Pfurtscheller et al. demonstrated that mu band desynchronization during sensory and motor tasks and beta band resynchronization after the movement can be detected through EEG recordings [42, 43]. They showed that the amount of muscle mass engaged in the movement affects post-movement beta resynchronization, reflecting the brain’s adaptation to varying motor demands, with larger beta increases observed after gross wrist movements compared to more localized finger and thumb movements. The rebound of beta rhythm in the motor cortex plays a key role in resetting cortical states after movements [44]. The analysis of the EEG signal at these bands could effectively predict hand motion trajectories, emphasizing their role in movement preparation and execution [45]. Changes in mu and beta amplitude of the EEG during upper limb movement have been shown to correlate with motor impairment and structural damage in subacute stroke patients [46].

The variations in motor performance observed during finger movement tasks can be explored through both state-based and trait-based analyses. State-based analyses examine fluctuations in performance within individuals, focusing on how transient factors such as attention, fatigue, or emotional state influence motor function. For instance, research has shown that focusing attention on specific aspects of finger-tapping movements disrupts their automaticity and decreases the coherence and speed of the movements, displaying that attentional demands can negatively affect motor performance [47]. Similarly, fatigue can slow motor responses and increase movement variability, reflecting the dynamic impact of temporary conditions on motor control [48]. Fast and slow trials could be compared based on EMG onset latency, as latency reflects the subject’s state of performance, even when trials are recorded consecutively on the same day. EMG onset latency offers a measure of motor execution efficiency, with faster trials indicating more efficient sensorimotor processing and slower trials suggesting momentary lapses. Analyzing this variability within the same session helps reveal the dynamic influence of transient factors on motor performance [49].

Trait-based analyses, on the other hand, compare performance differences between individuals and attribute these variations to inherent characteristics such as age, skill level, personality, or long-term neural adaptations. Studies have demonstrated that ageing is associated with declines in motor performance, as structural and functional brain changes lead to reduced grey matter volume and altered neural activation patterns [50]. Personality traits can influence the success of imagining hand movements. Extroverts had higher success rates in imagining right-hand movements, suggesting an interaction between personality traits and neural control during hand movement imagination tasks [51]. Such findings highlight the importance of understanding both transient (state-based) and stable (trait-based) factors when assessing motor performance and designing interventions to improve motor function.

On the other hand, EMG has been an invaluable tool for studying the neuromuscular aspects of finger movement tasks. EMG measures muscle activation patterns, providing insights into the coordination and timing of muscle contractions during movement execution. While some studies have demonstrated that EMG activity correlates with movement speed and force production, where faster movements are typically associated with higher EMG amplitudes due to greater motor unit recruitment [52], other studies suggest that the relationship between EMG magnitude and muscle force is not straightforward. Factors such as motor unit recruitment strategies, firing rates, muscle fibre properties, and the non-linear nature of EMG-force relationships complicate direct correlations [53].

In the context of movement latency, EMG has been used to pinpoint the onset of muscle activity, offering measurements of motor initiation relative to sensory cues and neural events [54]. Delays in muscle activation, particularly in patients with motor disorders, have been shown to contribute to prolonged movement latencies and impaired motor performance. However, only very few studies investigated the relationships between baseline or resting electrophysiological features and response latency. Notably, a study by Greenhouse et al. examined the relationship between resting corticospinal excitability, GABA levels in the motor cortex, and reaction time in healthy individuals [55]. The researchers found that individuals with higher resting corticospinal excitability had faster reaction times. Overall, the study highlighted how variations in resting and baseline neural activity can influence motor performance, specifically reaction times. However, while some aspects of the relationship between resting corticospinal excitability and reaction time have been explored, the connection between electrophysiological activity at rest and reaction time in healthy individuals remains under-explored, indicating a gap this study aims to address.

Despite the extensive research on finger movement tasks, there is a need for more integrated approaches that combine state-based and trait-based analyses with neural and muscular assessments such as EEG and EMG. Few studies have comprehensively examined the interaction between neural activity and muscle activation in motor performance. In this paper, by exploring how transient state-based factors influence sequential motor execution and comparing them with more stable, trait-based characteristics, we aim to gain a deeper understanding of motor variability both within and between subjects. Integrating simultaneous EEG and EMG data provides a more holistic view of the dynamic processes that govern motor control, which is crucial for developing targeted interventions in clinical and rehabilitative settings. This approach helps identifying the specific neural and muscular mechanisms and biomarkers underlying variability in motor function and guiding personalized strategies for improving motor performance in individuals with motor impairments or neurological disorders.

## 2. Methodology

### 2.1. Experiment and dataset description

In this study, we analyzed a publicly available dataset including EEG and EMG recordings during a stereotypical sequential finger-pinching task [56]. The experiment was conducted on 52 healthy subjects. It had imagery and executive phases. In this paper, we focused on the executive trials because we were interested in the simultaneous study of EEG and EMG. Each executive trial began with a 2-second fixation cross on a black screen, after which the subject was instructed to move either the left or right hand fingers based on the instruction presented on the screen, which was shown for 3 seconds. The sequential motor task consisted of touching the thumb with the index, middle, ring and small finger in that order, once at each trial. The subject waited for the presentation of the go cue and then started doing the movement, once, until the go cue disappeared from the monitor. Following each movement, there was a uniformly distributed random rest time, varying from 2.1 to 2.8 seconds until the beginning of the next trial. Each run consisted of 20 trials for each hand, randomly interleaved, and each subject completed one run. In the final shared dataset, the resting time for all the trials was cut to 2 seconds. The timeline of the trials and the movement is shown in Fig. 1.

**Figure 1:**
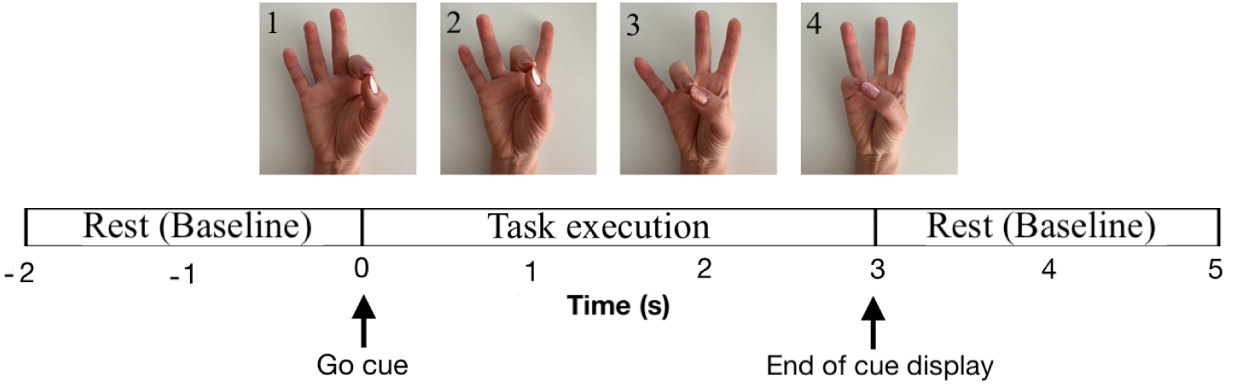
The finger-pinching task timeline. The top panel shows the finger movement sequence, where the subject pinches the thumb and each of the other fingers one by one. In the bottom panel, the timeline of the experiment is presented. The first two seconds were the rest time and the subject was waiting for the go cue display on a black screen. At *t* = 0 seconds the go cue was presented, indicating the hand to move, and the subject executed the corresponding movement. Then, the stimulus disappeared from the screen at *t* = 3 seconds and the subject rested, waiting for the next cue.

The Biosemi Active Two system with a 64-channel and BCI 2000 system was employed to record the EEG signals and present the instructions. EEG recordings were done following the international 10-10 system, with a sampling rate of 512 Hz. EMG signals were also recorded simultaneously using two electrodes attached to the flexor digitorum profundus and extensor digitorum of each arm to monitor muscle activity. The 3D coordinates of electrode locations were measured. The line noise was at 50 Hz.

### 2.2. Brain sources identification

To analyze the neural dynamics of the brain areas involved in the finger movement task, we needed to extract the corresponding brain sources from the EEG signals of each subject, cluster them based on selected features (as detailed in section 2.5), and then analyze the time series of each cluster to gain group-level insight into the relationship between the activity of related brain areas and subject performance. The initial steps involved cleaning and preprocessing the EEG and EMG signals. After cleaning the EEG data, we extracted the independent components (ICs) [57]. Examining the time series from EEG channels alone does not provide precise information about the activity within the sensorimotor area of the brain [58]. The signals recorded at each channel represent a mixture of activities from various brain circuits plus artifacts. Therefore, observing the activity at channels *C*3 and *C*4, which are positioned above the sensorimotor area, does not allow us to unequivocally attribute this activity to the sensorimotor area. This is because these channels also capture signals from adjacent brain regions. In this dataset, it was evident that for many subjects there was no discernible activity specifically originating from the sensorimotor area during the finger movement task. Thus, even after removing artifacts, the time series of these channels are not representative of the brain regions directly beneath them. Hence, we applied IC decomposition to make sure we only keep subjects with brain sources in sensorimotor areas.

To study the dynamics of the sensorimotor area, it is essential to decompose the EEG signals into ICs and spatially identify the specific brain regions associated with each IC. We accomplished this through automatic dipole fitting using EEGLAB (explained in subsections 2.4.1 to 2.4.3), enabling us to localize the sources of the extracted ICs [59]. A dipole fitted to an IC provides both the location and orientation of the neural population generating the time course of that IC. This method allows us to accurately attribute recorded activities to brain sources within the sensorimotor area.

### 2.3. Time reference

Since the exact timing of the finger movements was not recorded during the experiment, we needed to establish a time reference for each trial to measure subject performance. We identified the onset of the first finger movement— when the thumb and index finger touched— using the EMG signals from each trial. This EMG onset time was then used as the reference point for execution latency in the corresponding trial. The data preprocessing involved both EEG and EMG processing to extract brain dipole activity and EMG onset times, respectively.

2.4. *Cleaning and preprocessing the raw data*

The preprocessing of the recorded EEG and EMG signals was carried out in three main phases, as shown in the diagram in Fig. 2. The steps indicated in purple represent the processing applied to the combined EEG and EMG channels, while the steps indicated in blue and red steps were applied separately to the EEG and EMG signals, respectively. All preprocessing steps were performed using EEGLAB commands and extensions [60]. Unless otherwise specified, we used the default settings for these commands.

**Figure 2:**
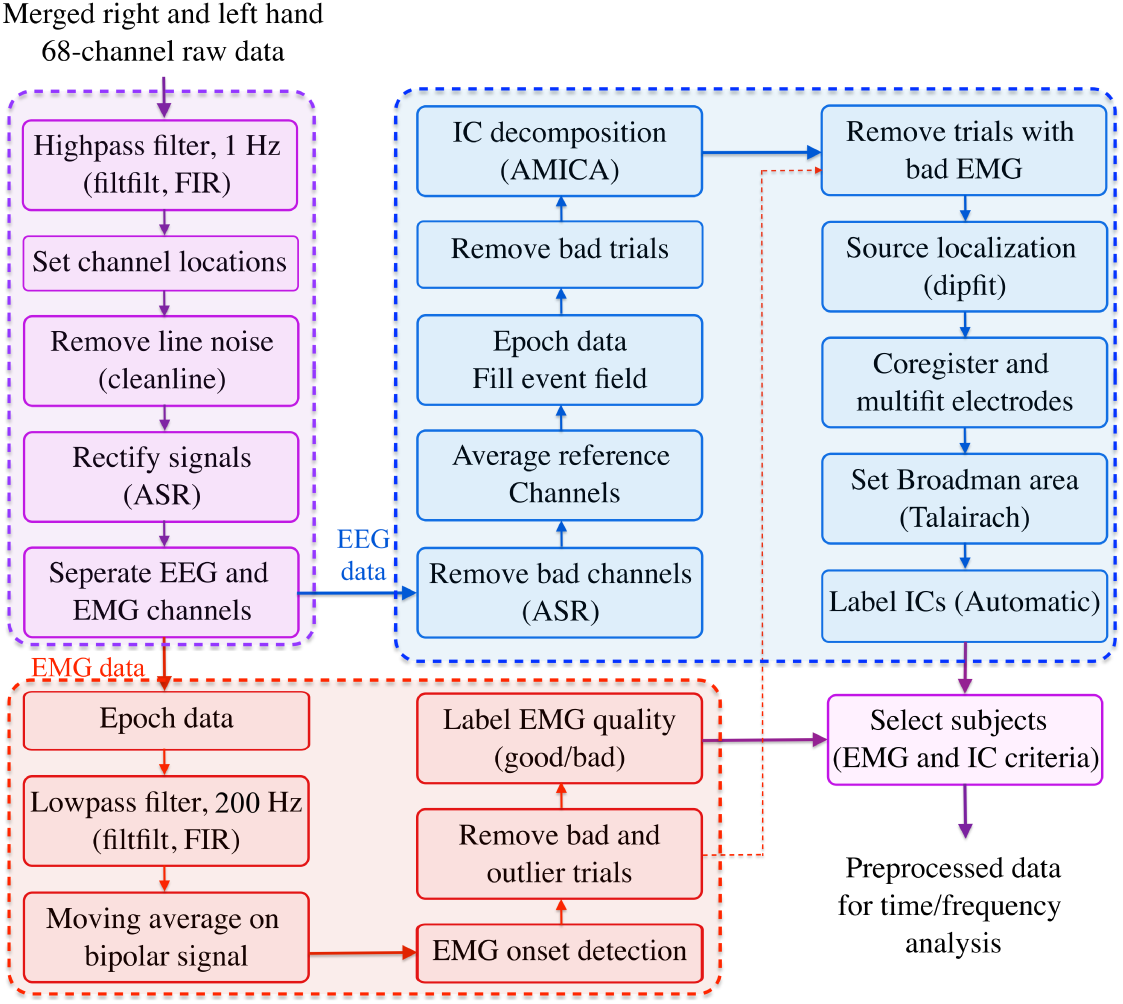
Preprocessing workflow: Processes shared between EEG and EMG signal preprocessing are highlighted in purple. Red blocks indicate procedures specific to EMG signal processing, while blue blocks denote those unique to EEG signal processing.

We combined the raw data from the right and the left hand trials to create a longer time series for IC decomposition. This approach was motivated by the need for extended time series data to enhance the accuracy of the IC decomposition process.

#### 2.4.1. Shared EEG and EMG signals preprocessing steps

The following steps were performed on the initial raw data from all 68 channels, including both EEG and EMG signals. We applied a high-pass filter at 1 Hz to the raw signals, a standard preprocessing step to improve the signal-to-noise ratio (SNR) and remove baseline drift. We used the cleanline extension of EEGLAB to remove 50 Hz line noise from the data. The data was processed using the artifact subspace reconstruction (ASR) extension of EEGLAB to address artifacts. Then, we separated the EMG and EEG channels for further cleaning and processing.

#### 2.4.2. EEG signals preprocessing

We then processed the 64 EEG channels’ data using the steps outlined below. Bad channels were removed using ASR with default parameters, except for the flat channel criterion, which was set to 4, and the minimum acceptable correlation with nearby channels, adjusted to 0.6 (instead of the default 0.8). These settings allow for the retention of more channels that may still contain valuable signal information despite slight deviations. Since the spatial resolution is crucial in studying the brain sources, keeping more channels by lowering the correlation threshold, helps to retain spatial information that would otherwise be lost if too many channels were removed. The EEG channels were re-referenced to the average, as the data were recorded with a common average reference. Then, the data was epoched into trials and bad trials were automatically removed. The signals were decomposed using Adaptive Mixture Independent Component Analysis (AMICA) [61]. Although AMICA requires more computational time than other decomposition methods, it has been shown to provide the most stable results [62, 63]. Trials with poor EMG recordings were excluded based on the preprocessing of the EMG data, which is detailed in the following subsection. Independent component sources were localized using the `dipfit` extension, applying the `pop dipfit settings` to identify the component sources. Then, EEG electrodes were coregistered, and the `multifit` function was used to align the electrodes with the head model and improve dipole localization. Then, BAs for the dipoles were identified [60]. Finally, the ICs were automatically labeled as either brain-derived or artifact.

#### 2.4.3. EMG signals preprocessing

The objective of cleaning the EMG data was to accurately determine the onset of finger movements for each trial, ensuring reliable and precise measurements for further analysis. The EMG preprocessing was done through the following steps.

The data was epoched into trials, as done for the EEG data. To remove noise, a low-pass filter was applied with a cutoff at 200 Hz. The signals from the two electrodes of the same hand were subtracted to obtain a bipolar signal, thus removing cardiac contributions from the EMG. This was followed by the application of a moving average filter to the bipolar signal with a window length of 15 samples, determined empirically. This filter reduced false EMG onset detections and missed true onsets. The EMG onset time was detected for each trial. To do so, first, the maximum amplitude of the bipolar EMG was identified after applying the moving average filter. A threshold was set to 25% of this maximum value, and the first instance where the signal crossed this threshold was marked as the onset time for the trial. Trials were discarded if the standard deviation (SD) of the bipolar EMG signal before the go cue exceeded one-third of the SD during the execution phase. Additionally, the interquartile range (IQR) of EMG onset times was computed for each subject. If the IQR was outside the range of [100 150] samples, it was set to the closest value in that range. The threshold for outlier detection was set to twice the IQR to accommodate variability in movement speed, as subjects were not instructed to perform the task at a specific speed. Subjects were labeled as ‘good’ or ‘bad’ based on trial discards: If more than 50% of a subject’s trials were discarded during preprocessing due to bad or outlier trials, the subject was labeled as ‘bad’. Otherwise, the subject was labeled as ‘good’ (for a detailed plot see Fig. 3 in [39]).

**Figure 3:**
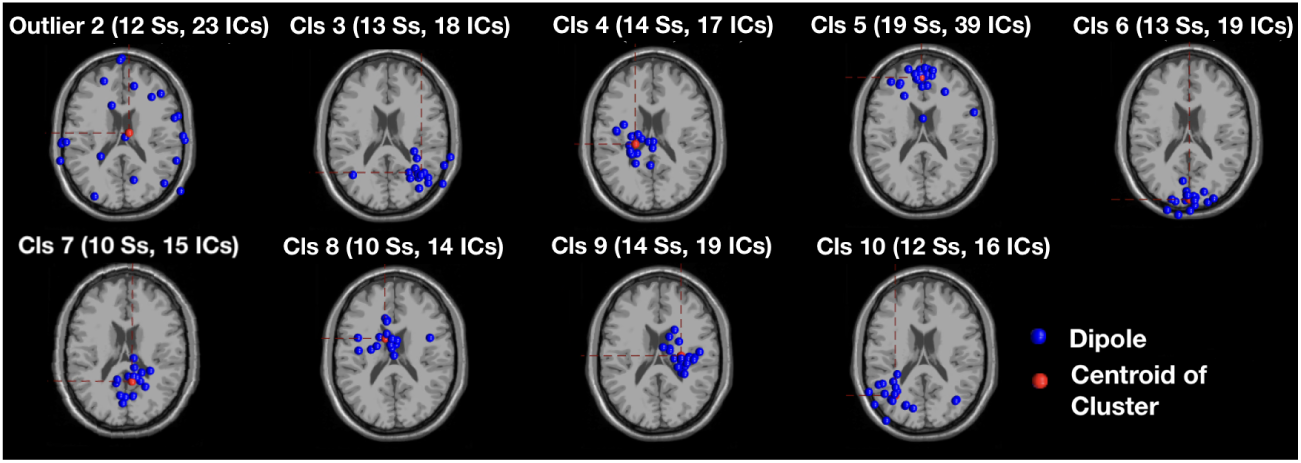
Initial clustering results: Clustering the dipoles into 8 clusters produced stable results, with consistent clustering patterns across multiple runs. A total of 19 subjects were included in the analysis. Notably, cluster 5 contains eye artifact components from all subjects. The corresponding scalp plot is shown for each cluster, with the number of subjects and ICs indicated above each plot. After cleaning, clusters 3 and 10 were selected as the right and left visual association areas and cluster 7 was selected as the visual area’s cluster. Clusters 4 and 9 were selected as the left and right sensorimotor clusters.

We established three criteria for including subjects in the further analysis. The first criterion was that subjects must have at least one dipole in sensorimotor areas, with a probability greater than 60% of being a brain dipole. Second, the dipole fit had to exhibit a residual variance of less than 15%. Lastly, the subject must have been labeled as ‘good’ after EMG processing. By applying these criteria, 21 out of the initial 38 subjects were retained. During the clustering process, two additional subjects were excluded because the power spectrum profiles of their dipoles did not align with the other dipoles in the selected clusters, which we will discuss in the next section.

The elimination of 50% of the subjects due to the absence of acceptable brain dipoles in the sensorimotor area in a motor movement task highlights the importance of decomposing EEG signals into ICs and identifying the Brodmann areas (BAs) of the dipoles. These subjects would not be eliminated if we had only focused on the channels’ time series and analyzed the electrodes located over sensorimotor areas, i.e. *C*3 and *C*4. This is particularly important for studies investigating spatiotemporal dynamics and neural activities. Decomposing EEG signals into independent components and localizing them in specific BAs plays a critical role in studying neural activity and its spatial dynamics, enabling more precise source identification and functional interpretation [64–66].

### 2.5. Clustering the dipoles

Clustering the brain dipoles allows us to identify and analyze common spatial and temporal patterns across subjects, enhancing our understanding of the underlying neural mechanisms. By grouping dipoles with similar characteristics, we can detect consistent patterns of neural activation, facilitating group-level analysis [67–69]. This method aggregates data, enabling robust and generalizable conclusions about neural processes [70]. Time-frequency analysis of clustered dipoles reveals dynamic changes in neural oscillations, providing insights into the temporal evolution of brain activity and consistent oscillatory patterns associated with specific brain regions and cognitive processes. Additionally, clustering improves the spatial localization of neural sources [71], leading to more accurate and interpretable results regarding the involved brain regions.

EEGLAB supports group-level analysis through its STUDY feature, using `std editset` function, which enables clustering and group-level comparisons of brain ICs [60, 72]. The STUDY framework in EEGLAB allows efficient aggregation of data across subjects, enabling robust statistical analyses and visual comparisons at a group level. For clustering purposes, a pre-clustering stage is required, which involves calculating various metrics such as power spectra, Event-Related Spectral Perturbation (ERSP), and scalp maps of ICs. These metrics are then utilized to perform clustering. This approach enables the grouping of ICs that share similar patterns across multiple subjects, thereby facilitating the analysis of common brain dynamics within the group. The selected parameters for pre-clustering computations in our analysis were power spectra, dipole locations, and scalp maps, with corresponding weights equal to 1, 10, and 1, respectively. This yielded the initial clusters (Fig. 3). After obtaining the initial clusters, we proceeded with the cleaning procedure. The final components in each cluster met the following criteria: *i*) there was at most one component from each subject in each cluster to avoid over-weighting any single subject, and *ii*) the power spectrum and scalp map of the subjects should be similar to the group’s average, as determined by visual inspection. If a subject had more than one IC in a given cluster, we kept the IC that most closely matched the cluster’s average features and removed the rest. Applying these rules, we selected five cleaned clusters for further analysis, see Fig. 4. For a cluster to be considered, it should contain at least 50% of the subjects. The selected clusters met this criterion, except the left visual association area, which was retained to serve as the contralateral equivalent of the right visual association cluster. The final selected clusters were as follows:

**Figure 4:**
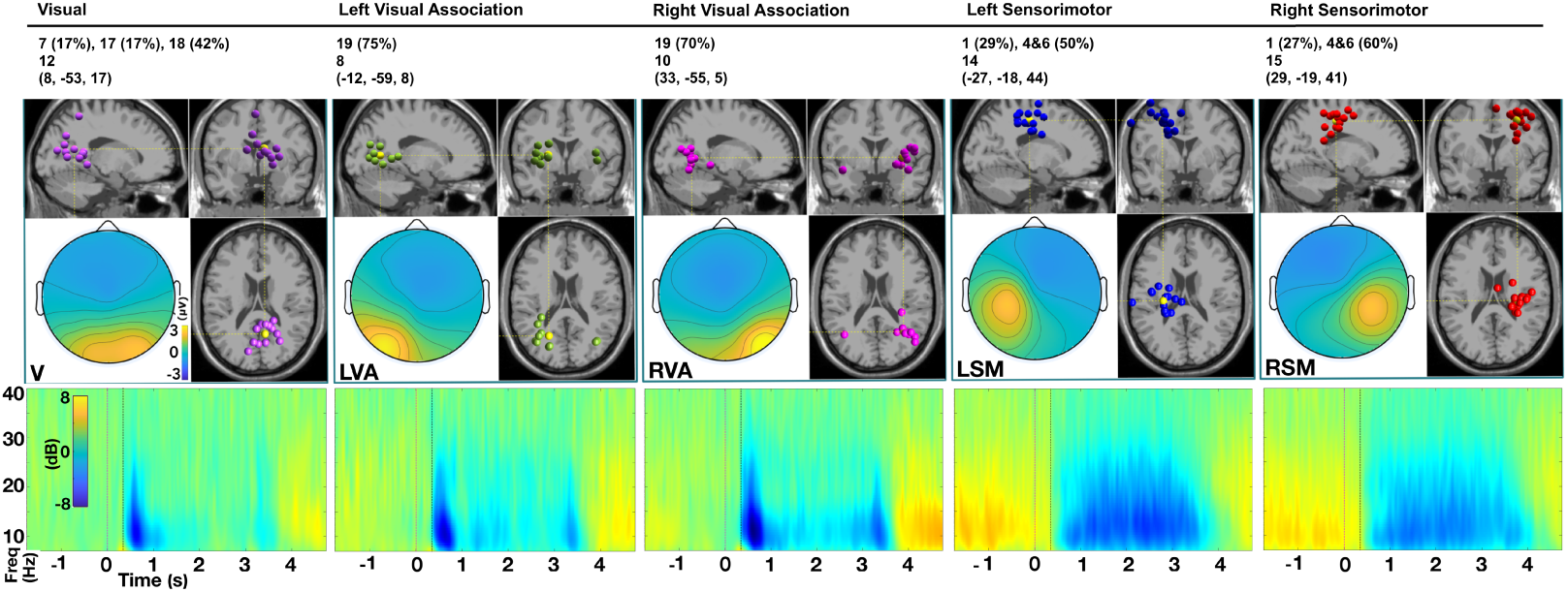
Clustered dipoles and corresponding ERSPs. The five selected clusters and their corresponding time-frequency power maps are displayed across five columns. The first row of the table on top shows, for each cluster, the percentage of dipoles that fit in the specified Brodmann area, the second row shows the number of contributing subjects, and the third row indicates the Talairach coordinates of the cluster centroid. In the first row of panels, the dipole locations for each cluster are shown in the sagittal, coronal, and horizontal planes of the brain. Additionally, a color-coded scalp map indicates the average topography for each cluster. The visual cluster is depicted in purple on the far left. The next two panels represent the left and right visual association areas, shown in green and magenta, respectively. The final two panels display the left and right sensorimotor areas, colored blue and red. The bottom row of panels illustrates the ERSPs for each cluster. The ERSPs for the first three clusters show a similar pattern: a desynchronization in the mu and low-beta bands following the stimulus, with a return to baseline after approximately 300 to 400 ms. The sensorimotor clusters display a decrease in the mu and low-beta bands as well, but with more delay, remaining below baseline until approximately 0.5 to 0.8 s after the end of the execution interval.

#### Visual (V) cluster

This cluster included 12 subjects, with dipoles located in the primary visual cortex (17%) and secondary visual cortex (42%), corresponding to BAs 17 and 18, respectively. BA 17 is heavily involved in retinotopic mapping, and BA 18 is involved in processing more complex visual stimuli, integrating information received from BA 17, and transmitting it to other higher-order visual areas. [73].

#### Left and Right Visual Association (LVA and RVA) clusters

These clusters contained 8 and 10 subjects, respectively. Where, 75% of the dipoles in LVA and 70% of them in RVA, belong to the BA 19, also known as the tertiary visual cortex, plays a critical role in higher-order visual processing. It integrates and interprets complex visual stimuli beyond basic features and recognizes complex patterns, motion, and visual object representation. It also integrates visual information through connections with multi-modal regions like the temporal and parietal lobes, contributing to the perception of movement and complex visual tasks [73, 74].

#### Left and Right sensorimotor (LSM and RSM) clusters

The LSM and RSM clusters included 14 and 15 subjects, respectively. In the LSM cluster, 29% of the dipoles were located in BA 1, which is part of the primary somatosensory cortex (SI), responsible for somatosensory processing [75], and 50% were in BA 4 and BA 6. In the RSM cluster, 27% of the dipoles were in BA 1, with 60% in BA 4 and BA 6. BA 4 corresponds to the primary motor cortex (M1), which is involved in motor execution and cognitive aspects like attention, motor learning, and movement inhibition. It also plays a role in integrating sensory information and modulating complex behaviors, critical for comprehensive motor control [76]. BA 6 encompasses both the premotor cortex and the supplementary motor area (SMA), which are involved in coordinating complex movements and integrating sensory input with motor commands, movement preparation and organization and cognitive integration [77].

### 2.6. Latency-based power analysis

After clustering the brain dipoles, and identifying their related brain regions, the next step was to classify the data based on the subjects’ executive latency. The goal was to explore the relationships between latency and neural activity in the selected brain regions and to determine the extent to which the differences between fast and slow responses reflect across-subjects (trait) or within-subjects (state) variability. To achieve this, we created two subsets of data for trait-based and state-based analysis.

#### Across-subjects analysis of traits

To assess individual performance traits, we conducted an across-subjects analysis focusing on response times. First, for each subject, we calculated the average response time by determining the mean of the EMG onset latencies across all trials. We then ranked the subjects according to their average response times. To classify the subjects, we defined fast performers as those whose average latencies were below the 30^th^ percentile and slow performers as those with latencies exceeding the 70^th^ percentile (Fig. 5). Finally, we pooled all individual trials from these two groups into separate categories: one representing the fast performers and another representing the slow performers.

**Figure 5:**
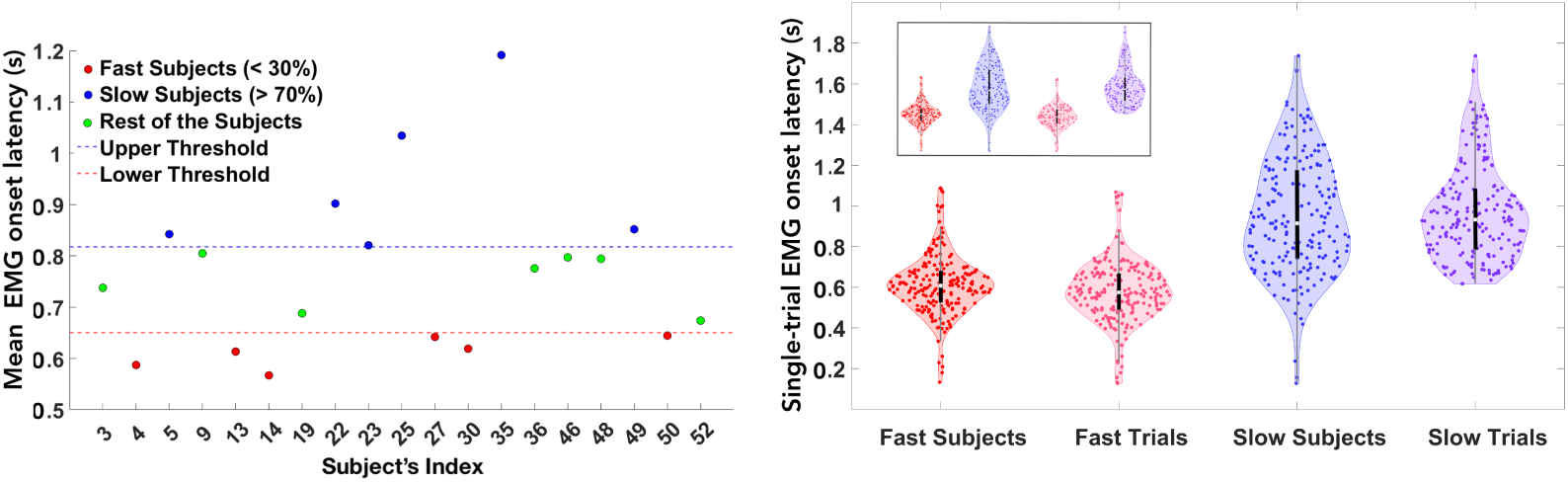
Trait and state-based latencies. Left: selection of fast and slow subjects based on average EMG onset latency, for trait-based analysis. Each dot represents the average EMG onset latency for an individual subject. The upper and lower thresholds for classification were set at 70^th^ and 30^th^ percentiles, respectively. Right: Violin plot of the EMG onset latencies for all the fast subjects, fast trials, slow subjects and slow trials, in red, pink, blue and purple color, respectively. Each dot inside the violins represents the EMG onset latency of a single trial in that subset. For each group, black circles indicate median values and vertical black lines show the interquartile range. The inset shows the trait and state-based EMG onset latencies side by side to provide more perspective.

#### Pooled within-subjects analysis of states

To examine within-subjects variations in performance across trials, we conducted a state analysis focusing on response times across individual trials. For each subject, we ranked the trials according to EMG onset latencies. We then classified the trials into two categories: fast trials, defined as those with latencies below the 30^th^ percentile, and slow trials, defined as those with latencies above the 70^th^ percentile. To assess neural dynamics related to response times, we pooled all fast trials from all subjects into a single group and did the same for the slow trials. This approach allowed us to compare the neural dynamics between the fast and slow trials at the group level, providing insights into latency-related changes across the entire subject pool.

The analysis of EMG onset latencies between fast and slow subjects and fast and slow trials reveals important distinctions in both trait-based and state-based muscle activation patterns (Table 1). Fast subjects showed a shorter mean EMG onset latency (0.61 s) compared to slow subjects (0.95 s), indicating that individuals classified as fast activate their muscles 340 ms earlier than their slow counterparts, on average. Additionally, fast subjects exhibited more consistent activation patterns, as seen in their lower variability (CV= 0.239) compared to slow subjects (CV= 0.300). This emphasizes the role of inherent subject characteristics, i.e., traits, in influencing muscle activation timing, where faster individuals have quicker and more reliable responses.

**Table 1:**
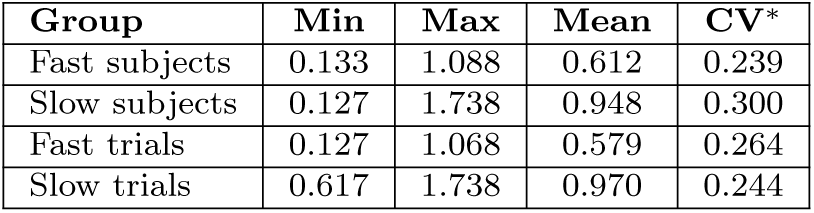
EMG onset time statistics: The minimum, maximum, mean and coefficient of variation (CV)^∗^ for the fast and slow subjects and trials.

| Group | Min | Max | Mean | CV* |
| --- | --- | --- | --- | --- |
| Fast subjects | 0.133 | 1.088 | 0.612 | 0.239 |
| Slow subjects | 0.127 | 1.738 | 0.948 | 0.300 |
| Fast trials | 0.127 | 1.068 | 0.579 | 0.264 |
| Slow trials | 0.617 | 1.738 | 0.970 | 0.244 |

In the state-based analysis, fast trials had a mean onset latency of 0.58 s, compared to 0.97 s in slow trials, with a gap of 390 ms. This suggests that the trial’s pace, irrespective of subject traits, had a stronger influence on muscle activation than inherent speed alone. The variability in fast trials was higher (CV= 0.264) than in slow trials (CV= 0.244). This difference was smaller than in the subject-based comparison, and contrary to that, here fast trials showed higher variation. All the trials of each participant were recorded within a single session, hence, the variability in performance defines the state-based differences. Since all trials were recorded in a single continuous session, state-based differences represent variability occurring at a relatively short timescale of up to 5 minutes, e.g., due to boredom or fluctuations in attention and arousal.

Next, we compared spectral power at each time point along the trial between fast and slow groups for both trait-based and state-based analyses. We focused on spectral power in the mu (7 to 14 Hz) and beta (15 to 30 Hz) bands, due to their relevance for motor behavior. This dual approach allowed us to explore EEG power variability addressing both between-subjects (trait) and within-subjects, between-trials (state) differences.

For the trait-based analysis, we concentrated on the between-subjects differences, emphasizing how consistent individual traits related to EEG power. We selected the fast and slow subjects based on their average EMG onset latencies (Fig. 5), which enabled us to identify distinct neural signatures associated with faster versus slower performers, revealing stable, trait-related differences in brain activity.

In contrast, in the state-based analysis, we focused on within-subjects variations across trials, emphasizing the differences in neural dynamics between fast and slow trials. This allowed us to study transient, latency-related changes in EEG power.

By combining the trait-based and state-based approaches, we aimed to gain a holistic understanding of the relationships between response time and EEG power. The between-subjects analysis provided insights into how consistent individual differences in latency (traits) were related to neural activity, while the within-subjects analysis highlighted how brain dynamics influence trial-level fluctuations (states).

2.7. *Balanced subsets for t-test analysis*

A common method for determining significant differences between subject groups or conditions in EEG data is the cluster-based permutation *t*-test, which handles the multiple comparisons problem effectively and is widely used in EEG/MEG analyses [78]. However, preprocessing and data cleaning, such as discarding bad trials, often result in an unequal number of trials per subject, even if the initial recordings were balanced. This introduces several challenges.

First, when aggregating trials at the subject level, subjects with more trials can dominate the results, causing uneven weighting across the dataset. Second, the standard *t*-test assumes homoscedasticity; the consequences of violating this assumption may be exacerbated by unequal trial counts, leading to inaccuracies in the results [79]. Additionally, EEG data often exhibit variability in data quality across subjects, introducing heterogeneity that a standard *t*-test does not account for [80].

To address these issues, we applied two corrective measures. First, to manage unequal trial counts, we resampled the data, ensuring that all subjects contributed an equal number of trials for both trait-based and state-based analyses. This eliminated the disproportionate influence of certain subjects. Table 2 shows the number of subjects contributing to the analyses for each brain area and each of the fast and slow groups in the state/trait-based comparisons. The #ST/IT columns show the number of trials after an equal number was selected for each subject (ST, selected trials), and the initial number of trials (IT, initial trials).

**Table 2:** Balanced data for statistical analysis: Number of subjects (Ss) and number of the selected trials over initial total trials (ST/IT) for the selected fast and slow groups, categorized by trait-based and state-based analyses across each brain area.

| Brain area | V |  | LVA |  | RVA |  | LSM |  | RSM |  |
| --- | --- | --- | --- | --- | --- | --- | --- | --- | --- | --- |
| Trait |  |  |  |  |  |  |  |  |  |  |
|  | #Ss | #ST/IT | #Ss | #ST/IT | #Ss | #ST/IT | #Ss | #ST/IT | #Ss | #ST/IT |
| Fast | 3 | 93/100 | 2 | 62/72 | 2 | 62/68 | 4 | 124/135 | 6 | 186/203 |
| Slow | 4 | 124/139 | 4 | 124/131 | 3 | 93/100 | 5 | 155/167 | 3 | 93/100 |
| State |  |  |  |  |  |  |  |  |  |  |
|  | #Ss | #ST/IT | #Ss | #ST/IT | #Ss | #ST/IT | #Ss | #ST/IT | #Ss | #ST/IT |
| Fast | 12 | 108/128 | 8 | 72/84 | 10 | 80/103 | 14 | 112/146 | 15 | 120/157 |
| Slow | 12 | 108/130 | 8 | 72/82 | 10 | 80/101 | 14 | 112/145 | 15 | 120/157 |

Second, to account for unequal variances, we used Welch’s *t*-test, [81]. This method is particularly suited for datasets with heteroscedasticity, improving the reliability of statistical comparisons when the assumption of equal variances is violated.

### 2.8. Spectral analyses and statistical tests

For both the fast and slow groups in the trait-based and state-based subsets, we performed a detailed time-frequency decomposition of the single trials across the ICs at each brain region of interest. The wavelet decomposition was carried out using the pop newtimef function from the EEGLAB toolbox, which provided a high-resolution time-frequency representation of the ICs activity at the single trial level. The average time-frequency decompositions of all the trials of all the ICs at each brain region are displayed in the bottom panel of Fig. 4 reflecting the average dynamic changes in brain oscillatory power in response to the go cue, indicating shared functional activity within the corresponding area. This allowed us to investigate the spectral dynamics associated with different response speeds. To specifically analyze the neural oscillatory activity, we focused on two frequency bands: the mu band (7 − 14 Hz) and the beta band (15 − 28 Hz). For each trial, we computed the average power within these bands, effectively capturing the brain’s rhythmic activity in regions related to motor control and sensory processing. We performed the same process on the bipolar EMG data, by computing the time-frequency decomposition of the single trials, and averaging over 20 to 200 Hz to extract the mean EMG power in this range for each trial.

Subsequently, we applied statistical testing to identify significant differences in power between the fast and slow groups. To this end, first, we ran a *t*-test for each time point over all the trials of the fast and slow groups for each brain area. Then, we employed a cluster-based permutation test, a robust non-parametric method ideal for EEG data where multiple comparisons are necessary. The test was performed with 1000 iterations. In each iteration, class labels were randomly shuffled, enabling us to build a null distribution of cluster sizes by computing the sum of the absolute value of the *t*-values of adjacent significant (at *p* = 0.05) time-points corresponding to the absence of differences between the two classes. Then, the empirical clusters obtained with the original class assignments were identified as significant clusters (SCs) at *p* = 0.01 or 0.05 if they exceeded the corresponding percentile of the null distribution. This method effectively handles the multiple comparisons problem by controlling for family-wise error rates, making it an appropriate choice for analyzing time series. The same approach was used for the EMG data, ensuring consistency in the analytical framework across both EEG and EMG modalities. We applied Welch’s *t*-test using the MATLAB function ttest2 with unequal variance in the two groups.

### 2.9. FDR-corrected results

In addition to the permutation tests, we performed a false discovery rate (FDR) correction to further validate our results and control for the possibility of false positives arising from multiple comparisons. Cluster-based permutation tests tend to sacrifice temporal precision because they group adjacent data points into clusters. The test assumes that nearby data points are correlated, i.e., not independent, and that significance should be considered in clusters rather than at isolated time points. This means that while the method is effective at controlling false positives, it can blur temporal boundaries, making it less precise in identifying the exact timing of significant effects [82]. On the other hand, FDR correction controls the proportion of false positives among the set of significant findings. It is more temporally precise compared to cluster-based methods because it evaluates the significance of each time point independently, without assuming clustering of effects. However, FDR can be more prone to false positives in noisy data compared to cluster-based methods, especially if the effects are weak or the signal is noisy. Also, it may miss extended temporal or spatial patterns of significance, as it treats each time point individually.

We conducted the FDR correction using the Benjamini-Hochberg procedure, implemented through the fdr bh function [83]. This method adjusts *p*-values to ensure that the expected proportion of false discoveries remains below a predefined threshold, which we set at *α* = 0.05. We reported FDR-corrected significant results at *p* = 0.05 and *p* = 0.01, as we did for the cluster-based tests. By controlling the FDR, we ensured that our results remained robust, even in the context of the large number of statistical tests conducted, thereby reducing the likelihood of spurious findings due to random fluctuations in the data.

By comparing the results of both methods, we could assess when the significant time points obtained with FDR were consistent with the significant clusters. Where both methods point to the same time intervals, we had higher confidence in the robustness and reliability of the findings.

Combining both FDR correction and cluster-based permutation tests offers a powerful way to balance temporal precision and robustness. While FDR correction ensures that we precisely identify the timing of significant effects, cluster-based permutation tests guard against false positives by identifying SCs over extended periods or regions. This combined approach is particularly useful for EEG/MEG studies where both exact timing and sustained effects are of interest. All the code for the preprocessing and analysis is available at: /github.com/GNB-UAM.

## 3. Results

After preprocessing the EEG and EMG data and conducting a group-level analysis to identify the brain sources associated with the finger-pinching task, we categorized the data into two subsets based on response latency to the go cue: a trait-based across-subjects subset and a state-based within-subjects subset. Each group comprised the 30% fastest and slowest responses, defined by the subject’s average latency for the trait-based category, and the single trial’s latency for the state-based category. We then applied a cluster-based permutation *t*-test and FDR correction to identify time intervals where the fast and slow groups exhibited significant differences in their band-limited time series, focusing on the mu and beta band power trajectories.

The outcomes of our analyses are represented in Figs. 6 to 8, which display the mean mu power, beta power, and broadband EMG power for both the fast and slow groups, along with their interquartile ranges (IQRs). These figures also highlight the clusters that exceeded the significance thresholds at *p* = 0.05 and 0.01 (herein referred to as SCs), as well as the significant time points at *p* = 0.05 and 0.01 as determined by the FDR correction for multiple comparisons.

**Figure 6:**
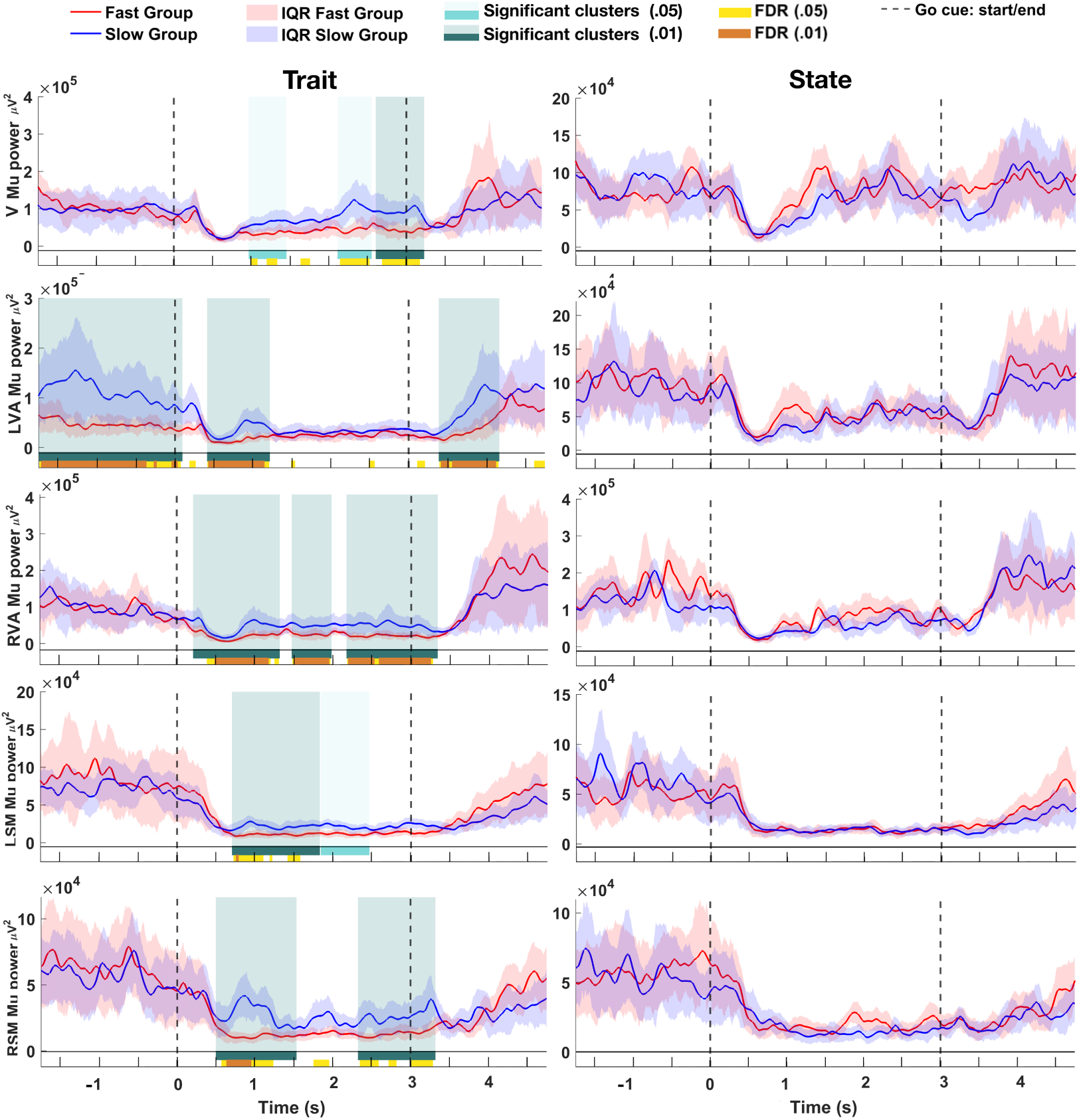
Response latency effects in the mu band are mostly observed during movement execution. The left column represents the fast and slow subjects (traits), and the right column corresponds to the fast and slow trials (states). Each row depicts one brain area, from top to bottom: V, LVA, RVA, LSM and RSM. The curves represent the average EEG power in the mu band across all single trials of the corresponding group, with shaded regions indicating the IQR (red, fast group; blue, slow group). The light and dark green shaded areas, along with matching colored rugs at the bottom of each plot, indicate intervals with SCs as identified by the permutation test, with *p*-value thresholds of 0.05 and 0.01, respectively. Additionally, yellow and orange rugs mark regions with significant *p*-values after FDR correction, with the same thresholds of 0.05 and 0.01, respectively.

**Figure 7:**
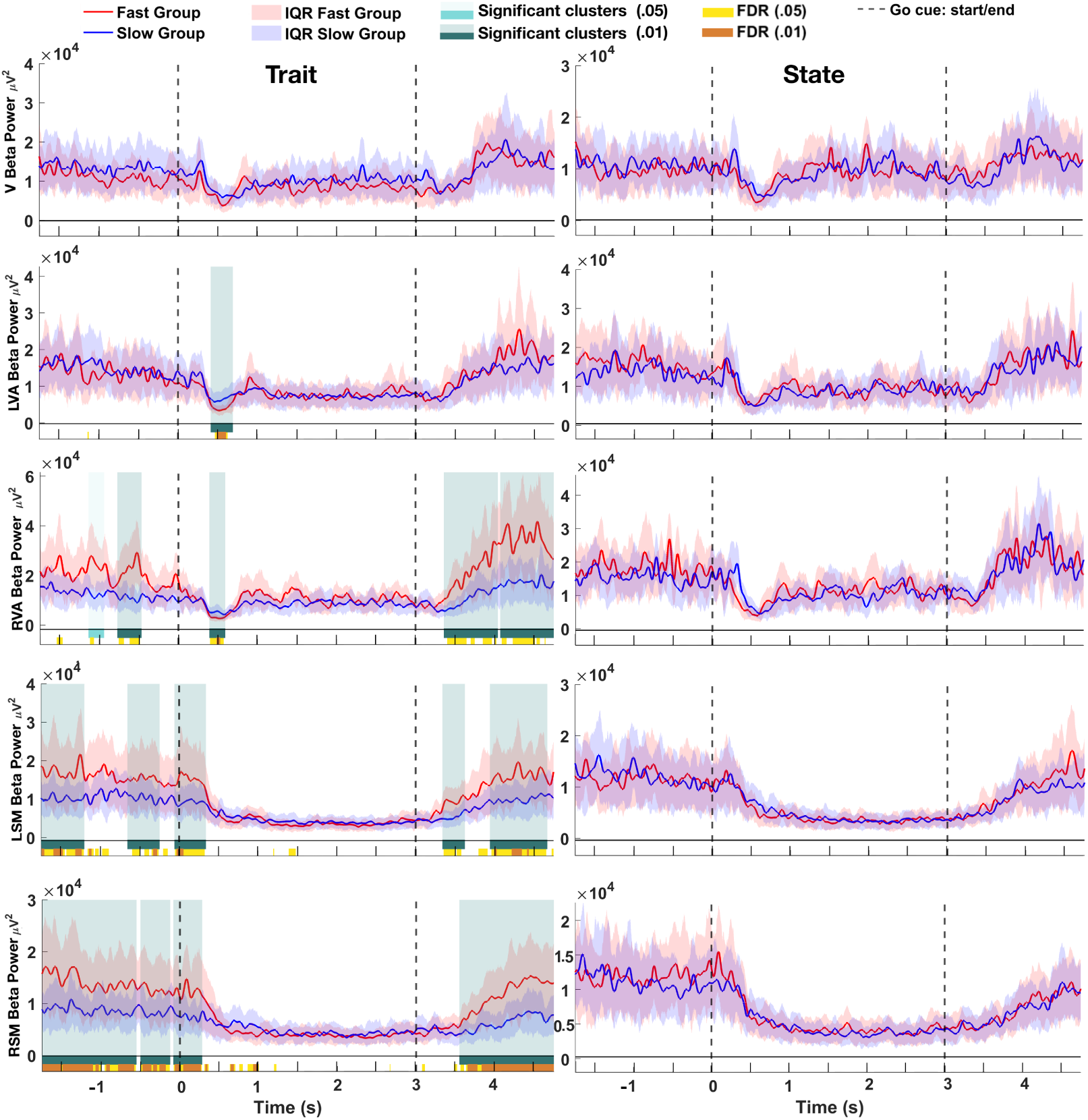
Response latency effects in the beta band are more prominent during the baseline period. The left column shows fast and slow subjects (traits), and the right column shows fast and slow trials (states). Rows represent brain areas: V, LVA, RVA, LSM, and RSM. Red and blue curves depict mean EEG power for fast and slow groups, respectively, with shaded areas indicating the IQR for each group. The light and dark green shaded regions, along with corresponding rugs, highlight SCs identified by the permutation test (*p*-values of 0.05 and 0.01). The yellow and orange rugs show FDR-corrected significant regions with the same thresholds of 0.05 and 0.01, respectively.

**Figure 8:**
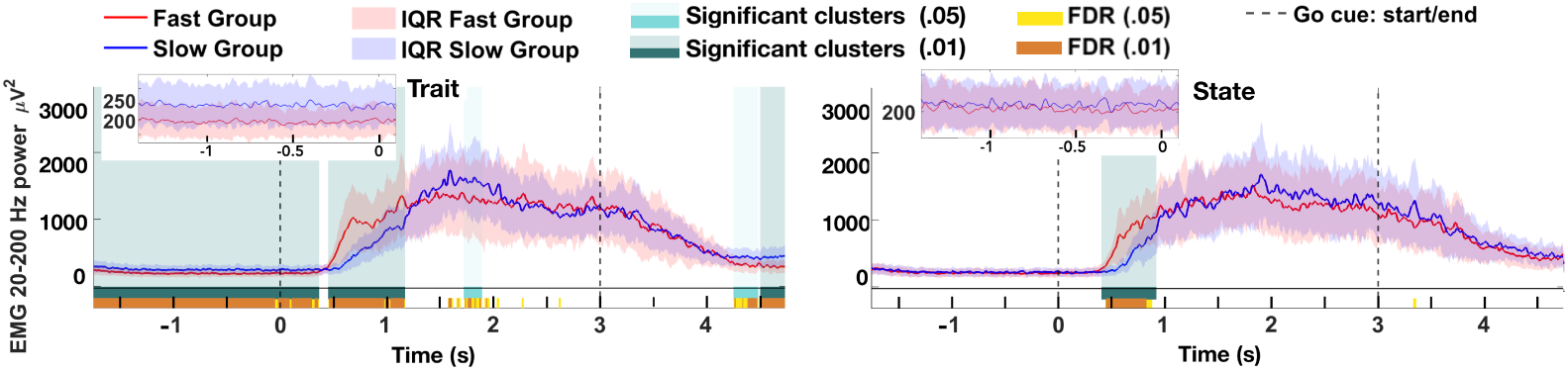
Fast responders exhibit lower EMG activity during the baseline period. The left plot displays fast and slow subjects, while the right plot shows fast and slow trials. The red and blue curves represent the mean EMG power for the fast and slow groups, with shaded regions showing the IQR for each group. The light and dark green shaded areas and corresponding rugs mark SCs detected by the permutation test (*p*-values= 0.05 and 0.01). Yellow and orange rugs indicate regions with FDR-corrected significance, using the same *p*-value thresholds.

The inspection of these figures provides a comprehensive view of how neural and muscular activity differ across subjects between fast and slow performers, as well as within-subjects between fast and slow trials.

### 3.1. Trait-based analysis

The trait-based analysis results showed significant intervals in all the brain areas in both the mu and beta bands, except the V area in the beta band. The fast and slow subjects showed notable differences in mu power across time and brain regions (see Fig. 6). SCs are indicated with shaded light green (*p* = 0.05) and dark green (*p* = 0.01) areas. These differences were more pronounced during the execution window in the mu band and the baseline period in the beta band. Below we inspect these results for each brain region and each frequency band in detail. First, we review the mu band results, as represented in Fig. 6.

#### V mu oscillations

Baseline: No significant difference was observed. Execution interval: moderately significant clusters (*p* = 0.05) emerged approximately 1 second after the go cue and again around 2.2 seconds after the go cue. A more significant cluster (*p* = 0.01) is observed near the end of the execution interval and could be related to processing visual feedback about the movement outcome, which is crucial for error detection and motor learning [84]. All three clusters contain FDR-corrected significant intervals highlighting the exact time of the significant differences between the two groups.

#### LVA mu oscillations

Baseline: Two SCs along with the FDR-corrected intervals were observed, corresponding to lower mu power in fast responders. This indicates sustained activity in the left visual association area in this subject group, suggesting heightened visual attention and readiness for the upcoming task. This anticipatory activation may facilitate quicker response times by priming the motor system [85, 86]. It also suggests involvement in evaluating the precision and success of the movement, possibly informing adjustments for future actions. Execution interval: An SC (with an FDR-corrected interval covering almost its full range) was detected starting around 300 ms after the go cue. This may reflect the processing of detailed visual information relevant to the task and coordination with motor commands [87].

#### RVA mu oscillations

Baseline: No significant difference was observed. Execution interval: Three substantial intervals of significant activity were observed, confirmed by FDR results. The first cluster started around 200 ms after the go cue and the last one ended approximately 200 ms after the movement interval, while the FDR approved the significant differences starting at ∼ 500 ms for *p* = 0.05, and around 600 ms for *p* = 0.01. These clusters were also significant at *p* = 0.01, indicating robust effects. The RVA’s robust activity points to its role in processing spatial and motion-related visual information essential for guiding movements [88, 89].

#### Sensorimotor mu oscillations

Baseline: No significant difference was observed. The lack of differences in this interval suggested that mu rhythms return to baseline similarly in both groups during non-movement periods, indicating that motor cortex activity is comparable between fast and slow responders outside of active movement. Execution interval: The fast group showed a more rapid and pronounced change in the mu band, characterized by higher and steeper desynchronization at the beginning of the movement. Mu rhythms are associated with the motor cortex at rest and are suppressed during movement execution [90]. The degree of mu suppression correlates with motor cortex activation levels [91]. Therefore, greater mu desynchronization in fast responders indicated more robust activation of motor areas during the task, facilitating quicker and more efficient movement execution, in line with previous studies [46].

The beta power differences between fast and slow subjects are observed across all brain regions except V. The rest of the brain areas show notable SCs, especially in the baseline and movement initiation phases, suggesting a stronger engagement in these regions for the fast group.

#### V beta oscillations

No SC was observed in the entire trial window.

#### LVA beta oscillations

Baseline: No SCs were detected. Execution interval: A SC (*p* = 0.01) began around 400 ms after the go cue, lasting approximately 250 ms; the FDR results show a later interval, starting around 500 ms after the go cue. Beta oscillations in visual areas may reflect the maintenance of the current visual state or the integration of visual information necessary for guiding motor actions [92, 93].

#### RVA beta oscillations

Two significant clusters (*p* = 0.05 and *p* = 0.01) were displayed starting at approximately 1150 and 700 ms before the go cue, lasting about 200 and 250 ms, respectively. Both clusters include FDR-corrected significant intervals (*p* = 0.05). The other cluster was observed starting about 350 ms after the end of the movement interval and extending until the end of the window. Elevated beta activity in the right visual association area during the baseline suggests that participants engaged in anticipatory visual processing in the non-executive interval. The right hemisphere is often associated with spatial attention and global visual processing [89]. High beta activity in the fast responders suggests involvement in processing visual feedback related to the movement. This may contribute to error detection and motor learning [84]. Execution interval: A SC was observed starting about 400 ms after the go cue, similar in timing to the one shown in the LVA.

#### Sensorimotor beta oscillations

Baseline: We observed SCs in this interval in both hemispheres where fast responders exhibited higher beta power. This reflects stronger maintenance of the current motor state and enhanced motor inhibition. Beta oscillations are associated with the suppression of movement and the maintenance of the status quo [92, 94]. Higher beta synchronization suggested that fast responders were better at inhibiting premature movements and more efficient in resetting the motor system, ensuring readiness without unnecessary motor activity. The post-movement beta rebound is thought to be involved in terminating the previous motor command and preparing the motor system for subsequent actions, contributing to faster response times in future tasks [95]. Execution interval: After the onset of the go cue, we still observed significant differences between the fast and slow groups. This cluster ended, right before SCs appeared in the LVA and RVA areas. The sequence of ending of LSM and RSM clusters and beginning of LVA and RVA’s SCs, respectively, manifested in cluster-based permutation test and FDR-corrected intervals. No significant differences were observed during the movement phase. Both fast and slow responders exhibited similarly low beta power during movement execution. Movement-related beta desynchronization reflects the release of motor inhibition to allow movement initiation, a process that yields similarly low beta power values during movement execution in the two groups but starts from different levels of baseline beta power, with fast responders exhibiting higher beta power in the precue window [96].

#### EMG activity

Baseline: We observed SCs along with FDR-corrected significant differences in the whole baseline window, starting around 1.3 s after the disappearance of the go cue. Lower EMG power in this interval suggests that fast responders had less muscle tension or pre-activation in the muscles involved in the finger-pinching task. This reduction indicates effective suppression of unnecessary muscle activity before the go cue, reflecting better motor inhibition at the muscular level [97]. Execution interval: Increased EMG power 500 ms after the go cue implies that fast responders generated an anticipated and stronger muscle contraction at the beginning of the movement. This suggests that fast responders quickly ramped up muscle activation, leading to faster and more forceful movements.

### 3.2. State-based analysis

The state-based results displayed no SC at any of the brain areas. This means that there were no consistent patterns of differences between the fast and slow trials in the considered frequency bands, suggesting that brain activity related to fast and slow trials was quite similar. The similar averages and overlapping IQRs indicated that the central tendencies and data distribution between the fast and slow trials were highly alike. The lack of significant differences implies that the fast and slow trials did not show distinct neural markers, suggesting that the speed of response might not be strongly tied to specific, observable changes in the EEG signals in this particular state-based analysis. It indicates that within-subjects variability in response times in this task is not driven by notable changes in neural activity in the mu and beta bands.

In the results presented in Fig. 6 to Fig. 8, we observed some FDR-corrected significant intervals that were not included in any SC. These intervals might reflect real, short-lived and temporally precise differences that did not meet the criteria to form a cluster in the permutation test. In motor or cognitive tasks, brief but highly localized effects could be meaningful, representing sharp transitions in neural dynamics that are important for processes like decision-making, motor planning, or execution [98–100]. Alternatively, the FDR-corrected intervals outside the cluster-based significant regions might represent false positives that fall within the acceptable false discovery rate. FDR correction controls the overall false discovery rate, hence isolated significant time points are more likely to be false positives than continuous intervals of significant time points. Given that FDR-corrected significance is designed to limit the proportion of false positives, these intervals should be interpreted cautiously, but not entirely dismissed.

## 4. Discussion

We investigated the neural and muscular mechanisms underlying motor responses during a sequential finger-pinching task by analyzing EEG and EMG data. Participants were categorized into fast and slow responders based on their reaction times, allowing for both trait-based and state-based analyses.

In the EEG trait-based analysis, we observed differences in the pattern of fast and slow subjects between the right and left visual association areas. While both hemispheres displayed SCs in response to the go cue, their baseline activity exhibited different dynamics in the mu and beta bands. The simultaneously significant mu and beta activity in both hemispheres shortly after the go cue suggests bilateral involvement of visual association areas in processing visual stimuli and integrating this information with motor planning. However, the presence of precue significant effects in mu power in the LVA (but not in RVA) and in beta power in the RVA (but not in the LVA) suggests possible hemispheric specializations. In the beta band, the right hemisphere of fast participants showed more prominent activity, indicating its greater involvement in processing visual feedback for motor tasks.

The observed SCs in BAs 7, 17, 18, and 19 during the different phases of the task suggest that these visual processing regions contribute to motor performance through anticipatory attention, visuomotor integration, and feedback processing. Anticipatory visual attention prepares the related circuits before movement, visuomotor integration facilitates coordination during the movement, and feedback processing aids in post-movement adjustments [85, 101–103]. The RVA’s association with spatial attention and global visual processing further supports its specialized role [89, 104]. Beta oscillations in these regions may indicate the maintenance of the current visual state and the suppression of irrelevant information, facilitating optimal readiness and efficient visual feedback processing [92]. The lateralization of the mu activity in the left hemisphere and beta activity in the right hemisphere points to potential hemispheric specialization in visual processing and attentional mechanisms [105].

The trait-based results of the sensorimotor areas indicated greater mu desynchronization during movement execution in both hemispheres in fast responders compared to slow ones. Mu rhythm suppression signifies the release of motor inhibition and the initiation of motor commands and is associated with increased cortical excitability and active motor processing [90, 106, 107]. No significant differences were found in the baseline of the mu band, indicating similar baseline motor cortex activity outside active movement in this frequency band.

On the other hand, fast responders exhibited significantly higher baseline beta power in both hemispheres in sensorimotor areas compared to slow responders. This elevated beta activity suggests enhanced motor inhibition that suppresses unnecessary muscle activation, maintaining the system in an optimal state of readiness before movement initiation, as well as efficient resetting of motor networks after movement completion [92, 108]. Following the movement execution interval, fast responders efficiently reset their motor system, as indicated by rebounding beta power and releasing EMG activity. This resetting involves reestablishing motor inhibition and preparing the neural circuitry for subsequent actions, highlighting the sequential transition between different functional brain states to support continuous performance. This difference did not appear during the movement phase.

We observed a sequential order of appearance of significant differences between the fast and slow groups in the sensorimotor and visual association areas. In the beta band, the end of the baseline SC in the sensorimotor areas coincides with the start of the SC in the visual association area. Also, in the mu band, the SC in the RVA and LVA appear before the SC in the RSM and LSM. This suggests a sequential structure in the dynamics of brain activities between sensorimotor and visual association regions in the mu and beta bands.

In the EMG trait-based analysis, the observation of the reduced baseline muscle activity in the fast responders implies effective motor inhibition at the muscular level, preventing premature activation and conserving energy. Additionally, the significantly higher EMG power during the initial phase of the movement reflects the effective translation of neural signals into muscular action, resulting in stronger, rapid and robust initial muscle activation in fast responders, as expected. This was aligned with the elevated baseline beta power in the sensorimotor areas of both hemispheres.

In the EEG state-based analysis, where fast and slow trials within all participants were compared, no significant differences were observed between fast and slow trials. This absence of significant differences at the cortical level suggests that neural activity patterns measured by EEG during these intervals are similar between fast and slow trials. It implies that the neural mechanisms contributing to faster responses may not be fully captured by state-based EEG analysis or may occur outside the analyzed frequency bands.

On the other hand, the state-based EMG analysis revealed significant differences between fast and slow trials over a short interval after movement initiation. This finding indicates that the primary distinctions in within-subjects motor performance are only manifested at the muscular level, and are not reflected in cortical mu and beta activity in our analyses. This reflects a more efficient translation of neural commands into muscle activity during the initial phase of movement execution in fast responses.

The observations from both EEG and EMG recordings provide a coherent picture of the neuromuscular strategies employed during fast responses. Enhanced baseline motor inhibition, evidenced by higher EEG beta power and lower baseline EMG activity, allows for optimal readiness and prevents premature muscle activation. Stronger motor cortex activation during movement execution, indicated by greater EEG mu desynchronization, coupled with increased and anticipated EMG power in the initial movement phase, corresponds to rapid and efficient motor responses.

We showed that fast responders exhibit greater decreases in power during movement (with respect to the precue window) than slow responders in both the mu and the beta band in motor areas. However, the time windows where significant differences occur differ: mu power differs during movement, being lower for fast responders, while beta power differs mostly in the baseline window. Notably, had we performed baseline normalization as commonly done in motor studies examining mu or beta event-related desynchronization, we could not have dissociated the differential effects of fast response trait on the two rhythms, particularly the different task phases when they occur.

To our knowledge, only a few studies examined EEG activity without baseline normalization, which can explain why precue power variability has received little attention so far. For example, Mazaheri et al. showed that higher levels of prestimulus alpha and mu activity predict a higher likelihood of failing to inhibit motor responses, indicating a connection between pre- cue brain rhythms and fluctuations in sensorimotor performance [109]. A study by Buchholz et al. explored the role of prestimulus alpha and beta oscillations in sensorimotor coordination using a tactile go cue to one of two non-visible index fingers and a crossed hands posture, hence implementing a degree of sensorimotor incongruence [110]. They reported a small but positive correlation between beta activity in somatosensory areas and saccadic reaction time. Espenhahn et al. found that older adults exhibited higher pre-movement baseline beta power than younger adults, even though their performance in a visuomotor learning task was comparable. However, their analysis was performed directly on the electrodes’ activity, without examining the brain sources [111]. Iwama et al. found that pre-movement beta rhythmicity in the motor cortex enhances neural coupling, which is associated with maintaining more stable coordination of finger movements during tasks [112].

Muralidharan and Aron examined how inducing a high beta frequency state in the sensorimotor cortex affects movement [113]. Using a behavioral paradigm designed to enhance beta oscillations, they found that an increased beta state in the sensorimotor cortex leads to a significant slowing of motor movements. In contrast to that and in accordance with our results, Leriche et al. explored how sensorimotor beta dynamics relate to movement states in healthy individuals [94]. Their study suggests that reduced beta activity in the sensorimotor cortex is associated with a “slowed movement state”, indicating that lower beta power might reflect a neural condition conducive to reduced motor speed or readiness. Our study along with [94] focuses on observing natural variations in beta power and how these correlate with movement states in a non-manipulative setup, examining spontaneous changes in beta activity without artificially altering or inducing beta states. In contrast, Muralidharan and Aron’s study involved behavioral induction of a high beta state by presenting a second go cue shortly after the end of the first movement—around the time of peak beta resynchronization—in 20% of the trials, creating an experimental condition to directly assess the effects of heightened beta activity on movement, thereby actively manipulating neural oscillations to study their impact. These studies seem contradictory at first: reduced beta power is associated with slow responses in our study along with Leriche et al., while increased beta power leads to movement slowing in Muralidharan and Aron. However, these differences can be explained by the context and dynamics of beta oscillations. Beta oscillations may serve different roles depending on the baseline state and the timing of modulation. Elevated beta due to post-movement rebound may have a multiphasic effect—inhibitory initially, but priming over time. In other words, a high beta level may have a delaying effect on the response latency of the next movement if it is happening with a small temporal distance (within ∼ 1 s) post-movement. On the other hand, a low beta level at the baseline—with enough temporal distance (⪆ 3 s) from the previous movement—suggests a lack of motor readiness. Furthermore, the relatively low probability of the second go cue in their study could also have affected their results, as post-movement beta rebound is also associated with uncertainty estimation and the update of internal models [114]. These processes and their associated cognitive load could have affected the relationship between beta power and movement readiness beyond what could be ascribed to a purely pre-movement effect. Together, these studies suggest that beta oscillations have complex, context-dependent roles in motor control, with both excessive suppression and insufficient activation impacting movement speed.

More generally, these partially conflicting results can be reconciled by considering that post-movement beta rebound and pre-movement beta might serve different purposes. The dataset we analyzed was already segmented and hence did not enable us to investigate the continuous dynamics of beta activity from the end of one trial to the start of the next. However, we anticipate that a detailed analysis of inter-movement intervals will yield important insights into the different roles of beta as subjects progress from the post-movement window of one trial to the pre-movement window of the next trial. While motor inhibition is similarly required in both the post-movement and pre-movement phases (albeit under different kinematics states), other roles of beta might be differentially expressed in these two phases. We can expect resetting of sensorimotor networks and adaptive feedback integration to occur shortly after movement [44, 95, 114], and motor preparation in the null space [115, 116], informed by temporal predictions [117, 118], to become more prominent as the go cue approaches. When the interval between two successive movements is unexpectedly reduced, as in Muralidharan and Aron’s study, post-movement and pre-movement beta might interact in complex and nonlinear ways. Furthermore, the specificity of each motor task, in particular with regard to its sequentiality and difficulty, might be reflected in distinct frequency band dynamics. Importantly, the separate trait and state analyses conducted here enabled us to attribute the electrophysiological correlates of fast responses to subject-specific traits, rather than within-subjects state fluctuations—a distinction that has been largely overlooked in previous studies.

Our approach is purely correlative and hence does not enable us to determine causal relationships. However, our results suggest that interventions aimed to enhance beta rhythmicity in the sensorimotor cortex could potentially improve motor performance. For example, transcranial alternating current stimulation (tACS) at beta frequency could engage plasticity mechanisms resulting in enhanced beta activity, potentially leading to improved sensorimotor function. Previous studies showing facilitation of motor inhibition [119] and motor learning [120] by beta tACS support this hypothesis.

## 5. Conclusion

This study offers new insights into the neuromuscular dynamics underlying motor responses in a latency-based analysis of a sequential finger-pinching task. Participants with low response latency exhibited a distinct sequence of neural events, suggesting that fast responders achieve quicker motor responses through a coordinated sequence of neural dynamics: effective neural inhibition in the prestimulus baseline, evidenced by higher sensorimotor beta power, higher RVA beta power and lower EMG activity, along with enhanced visual attention as reflected by lower LVA mu power; robust neural activation during movement, demonstrated by a greater mu desynchronization first at both LVA and RVA followed by LSM and RSM, and higher EMG power; and, finally, efficient neural resetting after movement completion, with higher baseline beta rebound in sensorimotor areas and RVA. The greater magnitude of the dynamic interplay between neural inhibition and activation can enhance their motor performance compared to slower responders. This was also reflected in the sequence of changes in the dynamics of the EMG signals. Overall, these findings enhance our understanding of the temporal coordination and neural dynamics underlying rapid motor actions. By emphasizing the sequential activities, from motor inhibition to activation and resetting, in both motor and visual cortical areas, we highlight the importance of timing and magnitude of neural oscillations in response latency as a trait. This understanding opens up possibilities for developing training and rehabilitation strategies that target these sequential neural processes to enhance motor function. Specifically, beta-band tACS offers a promising intervention, with the potential to confirm the role of baseline beta oscillations in motor response speed in healthy individuals, and to serve as a therapeutic tool for patients or aging populations experiencing motor impairments.

Understanding these neural dynamics could inform the development of more responsive and efficient BCI systems by incorporating the identified patterns of neural inhibition and activation to enhance BCI control strategies. Particularly, in the context of motor rehabilitation, a detailed characterization of motor response and its association with specific neural events and dynamics can lead to distinct treatment, and the development of novel rehabilitation benchmarking and biomarkers. Thus, the assessment of the evolution of the latency of the response during a motor rehabilitation process can yield personalized therapies. In this regard, subject-specific adaptations in BCIs and neuroprosthetics have proven to boost performance and contributed to the design of performance predictors [121–123].

Future research will be needed to confirm these findings in a larger population. The lack of video or kinematics recordings of the finger-pinches in this dataset, which prevents the assessment of actual movements and their precise timing, was a drawback that prevented us from diving deeper into unraveling the dynamical changes during these sequential tasks. Our study suggests the use of precise temporal references in this characterization. Despite these limitations, the methodology and results discussed in this paper provide promising insights for applications in the design of personalized adaptations and motor performance indicators in a wide variety of neurotechnological applications.

## Funding

This research was supported by grants PID2021-122347NB-I00, 01621-01493911, and CPP2023-010818 (MCIN/AEI and ERDF-“A way of making Europe”).

## Notes

### Competing Interest Statement

The authors have declared no competing interest.

